# Prediction-error signals to violated expectations about person identity and head orientation are doubly-dissociated across the dorsal and ventral visual streams

**DOI:** 10.1101/471276

**Authors:** Jonathan E. Robinson, Will Woods, Sumie Leung, Jordy Kaufman, Michael Breakspear, Andrew W. Young, Patrick J. Johnston

**Affiliations:** School of Psychology and Counselling, Queensland University of Technology, Victoria Park Road, Kelvin Grove, QLD 4059, Australia; Institute of Health and Biomedical Innovation, 60 Musk Avenue, Kelvin Grove, QLD 4059, Australia; QIMR Berghofer Medical Research Institute, 300 Herston Road, Herston, QLD 4006, Australia; Brain & Psychological Sciences Research Centre, Swinburne University of Technology, Burwood Road, Hawthorn, VIC 3123, Australia; Department of Psychology, University of York, Heslington, YO10 5DD, United Kingdom

## Abstract

Predictive coding theories of perception suggest the importance of constantly updated internal models of the world in predicting future sensory inputs. One implication of such models is that cortical regions whose function is to resolve particular stimulus attributes should also signal prediction violations with respect to those same stimulus attributes. Previously, through carefully designed experiments, we have demonstrated early-mid latency EEG/MEG prediction-error signals in the dorsal visual stream to violated expectations about stimulus orientation/trajectory, with localisations consistent with cortical areas processing motion and orientation. Here we extend those methods to simultaneously investigate the predictive processes in both dorsal and ventral visual streams. In this MEG study we employed a contextual trajectory paradigm that builds expectations using a series of image presentations. We created expectations about both face orientation and identity, either of which can subsequently be violated. Crucially this paradigm allows us to parametrically test double dissociations between these different types of violations. The study identified double dissociations across the type of violation in the dorsal and ventral visual streams, such that the right fusiform gyrus showed greater evidence of prediction-error signals to Identity violations than to Orientation violations, whereas the left angular gyrus and postcentral gyrus showed the opposite pattern of results. Our results suggest comparable processes for error checking and context updating in high-level expectations instantiated across both perceptual streams. Perceptual prediction-error signalling is initiated in regions associated with the processing of different stimulus properties.

**Significance Statement:** Visual processing occurs along ‘what’ and ‘where’ information streams that run, respectively along the ventral and dorsal surface of the posterior brain. Predictive coding models of perception imply prediction-error detection processes that are instantiated at the level where particular stimulus attributes are parsed. This implies that, for instance, when considering face stimuli, signals arising through violated expectations about the person identity of the stimulus should localise to the ventral stream, whereas signals arising through violated expectations about head orientation should localise to the dorsal stream. We test this in a magnetoencephalography source localisation study. The analysis confirmed that prediction-error signals to identity versus head-orientation occur with similar latency, but activate doubly-dissociated brain regions along ventral and dorsal processing streams.

## Introduction

Predictive coding formulations of perception stress the importance of hierarchical prediction-error checking mechanisms that allow the brain to test internally generated models of the world against actual sensory input (Friston and Kiebel, 2009; Hohwy, 2013). A largely untested implication of such models is that error detection processes with respect to particular stimulus attributes should occur at the level of the processing hierarchy at which those attributes are resolved (Friston, 2005). The visual perceptual system provides an excellent test bed for these ideas because it is well established that distinct visual processing streams subserve particular stimulus attributes (Mishkin et al., 1983; Milner and Goodale, 1998). Mishkin et al. (1983) first described two channels of visual information processing: object recognition processes localise to regions along the ventral surface of the brain; whilst the processing of visuo-spatial information is localised to brain regions along the dorsal surface. Indeed, this division has been particularly well reflected in the study of face perception (Haxby et al., 2000; Andrews and Ewbank, 2004). The ventrally located fusiform gyrus (FG) has been identified as a cortical regions that subserves the recognition of face stimuli (Grill-Spector et al., 2004). By comparison, time varying aspects of the face such as head motion, head orientation, and facial emotional, brain activity are commonly localised to more dorsally located regions, such as the middle temporal, superior temporal and angular gyri (Allison et al., 2000; O’Toole et al., 2002; Carlin et al., 2011; Baseler et al., 2012).

In a previous study (Johnston et al., 2017) we identified early latency expectancy violation responses (∼110-210ms) to unexpected head and body orientations. These responses were localised to dorsal visual stream areas middle temporal, superior temporal and angular gyri. In another such study we identified that this early latency responses also arises in the violation of visual expectation in relation to face identity (Johnston et al., 2016) typically considered to involve the ventral stream. These studies suggested that responses in this time window may reflect similar processes relating to the integration of top-down and bottom-up information (particularly the detecting of informational mismatches between these). Our previous studies (Johnston et al., 2016; Robinson et al., 2018) also report a consistent, midlatency response to violated prediction in both identity and orientation violations, thought to be involved in higher level perceptual processing.

Crucially, however, despite evidence of a consistent signalling process to violated expectation across these attributes, it remains to be seen whether the signalling associated with violated expectations has a differential localisation, based on the aspect of the stimulus in which the violation occurs. It is also unclear if equivalent signalling processes can be identified in the time course of each localisation. Therefore, the present magnetoencephalography (MEG) study extends beyond the work in previous studies and aims to establish whether expectancy violation signals can be demonstrated in both dorsal and ventral stream processes within the same paradigm. Crucially it investigates whether these signals can be differentially localised to their respective perceptual streams.

The investigation combines a novel source localisation and time course analysis to identify double dissociations to violated expectation associated with the facial identity or the orientation trajectory of head stimulus image. The study adapts the contextual trajectory paradigm described in our previous studies (Johnston et al., 2017; Robinson et al., 2018) to build and subsequently violate expectation relating to both of these stimulus attributes. It was predicted that there would be a double dissociation in signalling during the early-mid response latency. Specifically, it was predicted that: 1) violated expectations about stimulus trajectory should result in responses localised to the dorsal, but not in the ventral stream regions; and 2) violated prediction about stimulus identity should result in responses that localise to the ventral but not in the dorsal stream regions.

## Materials and Methods

Determining the localisation of these signalling processes requires the use of brain imaging techniques with both high temporal and high spatial resolution. Previous studies (Simpson et al., 2015; Johnston et al., 2017) have successfully used Beamforming analysis to localise signalling in the M170 time window. The study described below used MEG localisation analysis developed from these previous studies to enable localisation of signalling responses to violated expectations of both the identity and orientation of a stimulus.

### Participants

Data were collected for twenty-two participants all of whom self-reported as right handed (11 female). The age of participants ranged between 19-48 years (*M* = 26.05 years, *SD* = 7.35 years). Participants were recruited from staff and students at Swinburne University of Technology (SUT), all of whom gave their informed consent to participate. The study was approved by the Human Research Ethics Committees of Swinburne University of Technology (SUT) and Queensland University of Technology.

### Stimuli

Face stimuli offered a means of creating violations in both ventral and dorsal streams, as the dorsal and ventral pathways involved in face perception are considered critical in neural models (Haxby et al., 2000) and robustly demonstrated (Kanwisher et al., 1997; Hoffman and Haxby, 2000). Face stimuli are also well known to elicit strong responses in the M170 time window (Eimer, 2011), that should maximise detection of responses to expectancy violations. Images of five exemplar faces were captured using a Professional 3D Graphics Rendering software (Poser 11) on a black background, at 2-degree orientation increments ranging between −26° and 34°, with 0° being the image face directly forward. Whilst it is usually desirable to use real faces, in this instance virtually rendered faces were used because of the difficulty of maintaining precise control over real faces in terms of changing only angle of orientation, whilst maintaining consistent eye gaze, light control, and facial expression. A constraint on maximal head orientation was defined because, unlike body stimuli, it is important that the face parts are not obscured in any image.

The range of orientations also needed to be wide enough to allow the creation of appropriate stimulus presentation sequences that could interrogate the factors of interest, whilst also maintaining optimal low-level stimulus controls. All exemplars were male, so that the space filled by hair and the shape of the face could be most easily maintained across exemplars (see Figure 1). Images were monochromatic and enclosed in a 300 × 300 pixel frame (72 ppi) using Adobe Photoshop (CC 2015) to mitigate lag in image loading during the experiment. Additionally, using MATLAB (v.2016b, MathWorks) for image adaptation, all exemplars were matched for approximate mean luminance (RGB pixel Value 117 +/− 1%) across each image set and resized to give them approximately similar black space (+/− 2%) (i.e. the area of black background not filled by the image subject). Duplicates of each image set were created with red dots added to each image in a random position, about the image subject. Any of these images could be used in the red dot vigilance task described in the procedure below.

**Figure 1.**
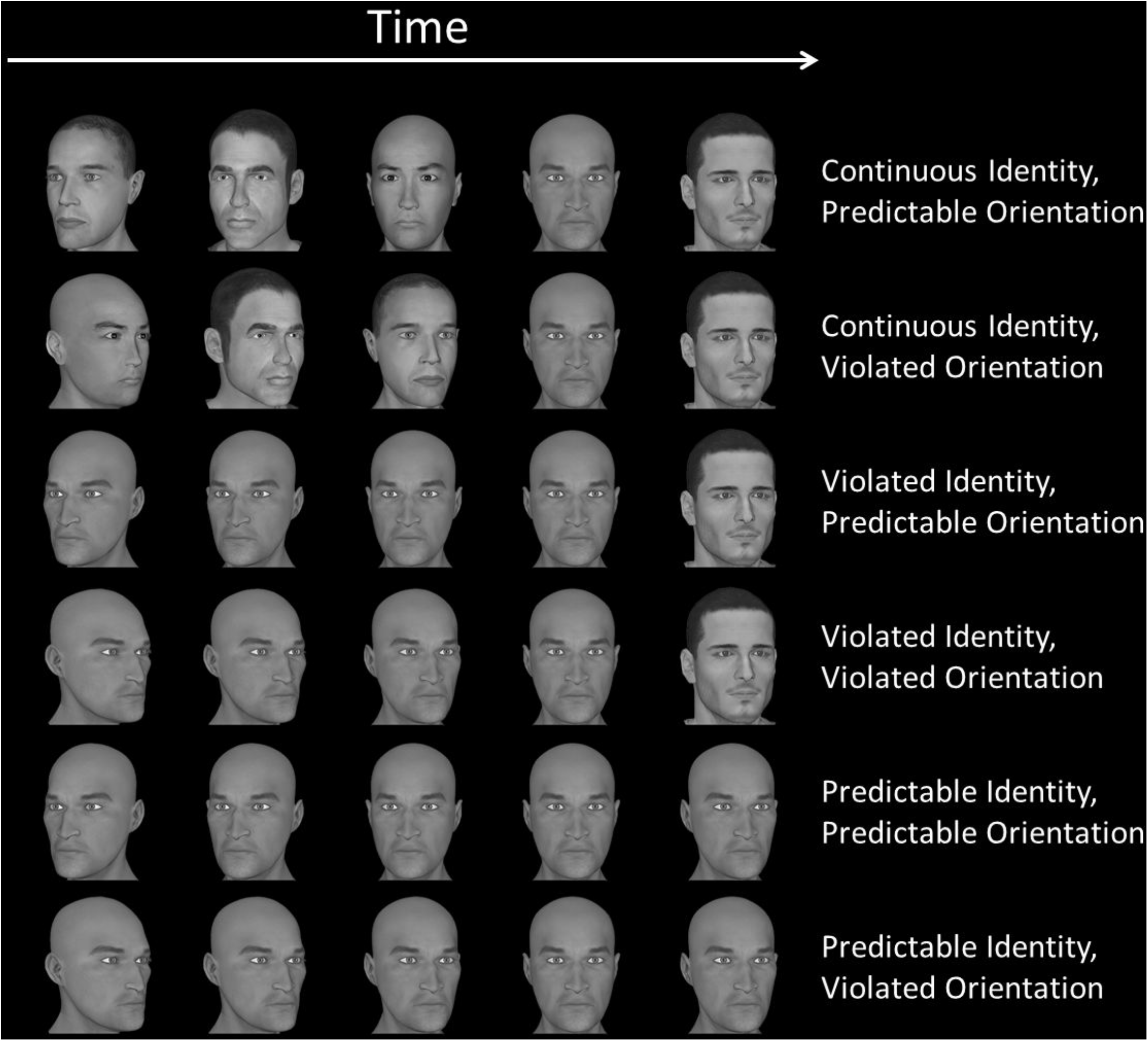
An example sequence for each of the six experimental conditions. With each pair (from top to bottom) representing the two Identity conditions (CCI, VI) and the control identity condition (PI), every other sequence (from top to bottom) represents the two possible Orientations (PO, VO). Note that each trial sequence across the four main conditions is matched to the final transition, and the two baseline conditions are also matched across the final transition.

Stimuli were presented using PsychoPy (v1.85.1) (Peirce, 2007, 2009) on a computer (Dell) activated by the experimenter and projected (Panasonic PT – D5100 projector) using a mirror periscope in the wall of the magnetically shielded room that surrounded the MEG. The periscope back projected the experiment onto a nonferrous projector screen (1.5 × 1 metres) placed approximately 1 metre from the participant, such that the stimuli filled approximately 2 degrees of the visual angle. A response box was also given to the participants so they could respond to catch trials.

### Experimental Design

The study used an adaptation of the contextual trajectory paradigm used in previous studies (Johnston et al., 2017; Robinson et al., 2018) to attempt to establish a double dissociation between expectancy violations relating to person identity and those relating head orientation trajectories. This study used a 2 × 2 repeated measures design—with factors of Expectation Type (Identity; Orientation) and Predictability (Non-violation; Violation). The individual conditions were, for the Identity Expectation Type: Continuously Changing Identity (CCI); Violated Identity (VI); and Orientation Expectation Type: Predictable Orientation (PO); Violated Orientation (VO). An additional control condition used a consistent identity with each Orientation: Predictable Identity Predictable Orientation (PIPO); and Predictable Identity Violated Orientation (PIVO).

### Procedure

Participants viewed sequences of five successive images of heads in each trial. Depending on the trial type, the final image in relation to the preceding sequence varied, to form different conditions. The first four images in each sequence established a clear trajectory of heads to create an expectation about orientation, and, except in CCI conditions, about person identity. The fifth and final image in each sequence either confirmed or violated the expectation about the Orientation or about the person’s Identity (see Figure 1). In the CCI conditions, all five images featured different character identities, so that no clear expectation about identity was established, but an expectation about orientation was established according to the paradigm.

As with the contextual trajectory paradigm used in Johnston et al. (2017), the design of the present experiment ensured a match in the final transition across Predictable Orientation (PO) and Violated Orientation (VO) conditions. This design also ensure that no trial sequence contained any identical stimulus repeat. This was achieved using multiple image transition increments at different head orientations and creating a matrix of sequences in each possible trajectory direction. The present experiment adapted this procedure, using multiple trials with two different orientation increments instead of three (8° and 10°), rotating in either a clockwise or anti-clockwise trajectory. These could then be collapsed across during the analysis.

In order to create identical final transitions across all conditions, it was important to include identity sequences that did not create surprise but had identically matched final transitions for both Identity Violated and non-violated Identity conditions. For this reason, the study used a continuously changing identity (CCI) condition (see Figure 1), where person identity changed in each step of the sequence, creating a lack of expectation person identity in the final transition and low surprise. By comparison, the Violated Identity (VI) trials presented a consistent identity for four stimuli with a changed identity presented in the final transition, Violating the accumulated expectation. Using this trial design, coupled with Orientation conditions used previously, these sequences form a set of trials that end with an identical final transition across both factors (Identity and Orientation). Because of these stringent controls, any differences in the ERF across these conditions must be attributable to the preceding context rather than any low-level stimulus differences between the final image onsets (because there are none).

It was important to ensure that not all sequences containing repeated identities were Violated identity trials. If this were the case, a Violated Identity could be assumed by the onset of the second image in the sequence and it’s outcome might become predictable. Therefore, as can be seen in Figure 1, two additional conditions (PIPO and PIVO), which replicate previous contextual trajectory paradigms (Johnston et al., 2017; Robinson et al., 2018) where a consistent or predictable identity (PI) was presented across all transitions of a trial and only orientation was manipulated. Each identity used appeared in all sequence positions, and all conditions, but contained a match across the four main experimental conditions (CCIPO, CCIVO, VIPO, VIVO) and separately across both of the control conditions (PIPO, PIVO). Furthermore, all of the sequences consistently used a penultimate image at a fixed transition point (4°) that enabled a match of stimuli in Orientation conditions. This transition point was intentionally offset from the centre (0°) to prevent the disruption of participants responding to known effects of being looked at (Jenkins et al., 2006). To maintain consistent likelihood of Violated Orientation half of all trials used a Predictable Orientation sequence and half used Violated Orientation sequence.

Stimuli were presented for 500ms, creating a total trial length (five stimuli) of 2.5 seconds, with a 500ms fixation cross presentation between each trial. To give the study sufficient power, 80 trials were presented randomly for each condition in the experiment, forming a total of 480 trials for analysis. In order to maintain participant vigilance throughout the experiment, an additional 60 “catch trials” were added, distributed randomly through the experiment (∼11% of the total trials). These trials featured a red dot images at a random point of the sequence, which participants were instructed to respond via a button press (using the button box provided) when they saw a red dot image. These catch trials were excluded from the ERF averaging. This gave the experiment a total of 540 trials, lasting approximately 27 minutes.

### MEG and MRI Acquisition Parameters

Data were acquired using the Elekta TRIUX MEG (306 channel; 102 magnetometers and 204 gradiometers) at the Brain and Psychological Sciences Centre (BPsyC), SUT. The data were recorded at a sample of 1000Hz, with an online high-pass filter of 0.1Hz and an online low-pass filter of 300Hz. Fiduciary electrodes were attached to both mastoids, and the left, right, and centre of the forehead in order to monitor the participants’ motion in the scanner. Electrodes were also attached above and below the right eye to monitor blinks, to the right wrist to monitor cardiac rhythm, and to the right elbow for grounding. The registration of the scalp in 3D space was achieved using a Polhemus 3D pen, which marked the position of each fiduciary electrode, the nasion, and both the left and right tragus, the pen was then used to draw the entire head shape in the 3D tracking software. The position of fiduciary electrodes was then recorded throughout the scan to ensure the head position could be tracked in relation to the sensors in the scan. This information and the 3D plot of the head shape could then be used later for coregistration with MRI scans. Participants were then tested for potential sources of noise in the magnetically silent scanner room, before being prepared for the acquisition. Once in the scanning room the participant was raised into the appropriate position under the sensor array and held in place using foam shims.

To enable anatomical inference in source-space analysis, each individual’s digitised head shape was coregistered with an individual anatomical MRI scan using surface matching (Kozinska et al., 2001). A high resolution T1-weighted structural MRI was acquired with a Siemens Tim Trio 3T MRI scanner, using a 12-channel head coil. The spatial resolution of the scan was 1.0 × 1.0 × 1.0mm, reconstructed to 1 mm isotropic voxels; a TR of 1900ms; a TE of 2520ms; a TI of 900ms; a flip-angle of 9°; and field-of-view of 256mm on a voxel matrix of 256 by 176. The total scan time for the T1 structural MRI was approximately 8 minutes. The MRI for each participant was segmented using FreeSurfer (v6.0.0) (Fischl et al., 2002), and a nonlinear transformation to the MNI 152 standard brain calculated using ANTS (Avants et al., 2009). A regular 5mm × 5mm × 5mm grid was defined on the MNI brain, and the inverse transformation applied to the grid points for each individual, resulting in a one-to-one anatomical correspondence for each grid point across the group.

### MEG Pre-processing

For each participant, MEG data were initially segmented into 3000ms windows around the final stimulus of each experimental trial, extending −2500ms prior to final stimulus onset, to encompass the entire trial, and 500ms after the final stimulus, in order to encompass the poststimulus response. These data segments were then manually inspected using a custom Python data visualisation application that visualised data variance simultaneously by trial and sensors. Sensors or trials assessed by the research team as outliers in the data, containing particularly high variance, were then excluded from further analysis. The data from the remaining magnetometers and gradiometers was filtered using a Butterworth filter using a high-pass of 1Hz and low-pass of 40Hz, with a slope −24 dB/octave.

### Statistical Analyses

In order to exploit the concurrent high spatial and temporal resolution of MEG, a two stage analysis strategy was applied. In the first stage spatial beamforming techniques were used to localise cortical regions that showed significant differences in activation to Predictable versus Violated Identity, and Predictable versus Violated Orientation. Identity comparisons used compound conditions of all Continuously Changing Identity (CCIPO, CCIVO) compared to all Violated Identity conditions (VIPO, VIVO). Orientation comparisons combined all conditions containing Predictable Orientations (CCIPO, VIPO, PIPO) and compared them to all conditions containing Violated Orientations (CCIVO, VIVO, PIVO). In stage 2, the virtual electrode time series for the optimum direction at each of the grid points at the cortical locations identified in stage 1 were analysed to identify *when* in these cortical locations the experimental conditions of interest showed statistically significant differences.

### Stage One: Localising Evoked Responses in Violated Identities and Violated Orientations using an Evoked Power Variance Metric in Spatial Beamformer Analysis

#### Overview

In order to identify brain regions that respond to Identity or Orientation Type Violations a beamforming analysis was performed. Here, however, this first step of the analysis aimed to localise activity associated with unexpected face Identities and unexpected head Orientation, not present in predictable conditions. This was achieved using two separate comparisons as described above. For each comparison the difference in the total power of the average evoked response for each condition was calculated for every location in source-space within the brain. In addition, a leave-one-out jackknife procedure was used to estimate the standard error of this difference at each location. The beamformer generates virtual electrode time series in source space for each epoch of data in the conditions being compared. These virtual electrodes invert signals from the sensors to model the time series at each location in a volumetric grid over the brain. To identify statistically significant local maxima at the group level a non-parametric permutation method was used.

#### Details of implementation

The beamformer requires the data covariance matrix ***C**_r_* which is calculated once, from all the data of both conditions to be contrasted during the active window. The active window is considered as 60ms to 500ms, where 0ms is the time of the presentation of the fifth image in each sequence. The leadfields used in the beamformer were calculated from a boundary element model of the inner skull using MNE-python (Gramfort et al., 2013; Gramfort et al., 2014). The beamformer weights at some position ***k*** with orientation ***q**_k_* are calculated:

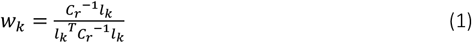

where

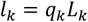

Here ***W**_k_* is the weight vector, ***L**_k_* is the three-dimensional leadfield at the point *k*, and ***C**_r_*, are the regularised estimates of covariance of the data for both conditions of interest combined; ***l**_k_* is the resultant 1D vector leadfield for the direction ***q**_k_*. For every point ***k*** on the volumetric grid, the resultant leadfield, and hence the weights ***w**_k_* are calculated for each of 150 approximately equally spaced orientations in a hemisphere derived using a Fibonacci lattice. The orientation vector ***q**_k_* is chosen to give the maximum difference in evoked power at each location *k* (using equation 4), in a similar manner to the Maximum Contrast minimum variance Beamformer introduced by Chen et al. (2006). In this case, however, no closed form solution for the optimal direction is available. For each of these 150 orientations, the dc corrected time series (using the period −200ms to 0ms) are passed through the weights to give the time series ***X_k,i_*(*t*), (*i* = 1,…, *N_X_*)** for that point and orientation

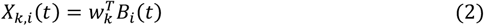

where ***β_i_*(*t*)** are the measurements from the sensors for each of the *i* presentations of a stimulus. The total power in the average evoked response for one condition is

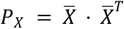

where 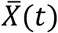 is the mean of the set of ***X_l,i_*** time series. The standard error of ***P_X_*** can be estimated using a leave-one-out jackknife procedure, which generates a new set ***P_Xjackknife_* = {*P_X1_, P_χ2_*,…, *P_XN_*}** where each of the ***P_Xj_*** are calculated using all the data except the j-th epoch. Then the standard error for ***P_X_*** can be estimated as

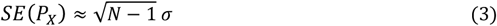

where ***ρ*** is the standard deviation of the jackknife samples ***P_Xjackknife_***.

The same calculation is applied to the second condition to give ***P_Y_***, ***SE*(*P_Y_*)** and the difference in evoked power calculated as a T-statistic

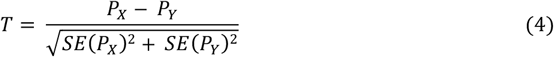

The largest ***T*** is chosen from all computed directions to represent the power difference at that point. This T-statistic was then constructed for every grid point in the brain, to give T-values across the whole brain for the comparison of interest. A group level statistical map was calculated as a one-sample T-statistic from the mean and standard error of the mean of the T values for each individual for each grid point. To test for statistically significant group differences, a non-parametric permutation procedure using maximum statistics to correct for the FWER was applied (Nichols and Holmes, 2002), under the null hypothesis that the condition labels X and Y are arbitrary, using only grid points included in a grey matter mask generated using the Harvard-Oxford cortical structural atlas parcellation (Frazier et al., 2005; Desikan et al., 2006; Makris et al., 2006; Goldstein et al., 2007) as implemented in the FSL (FMRIB, Software Library v5.0). This mask meant that only areas of the brain containing cortical grey matter were considered in the analysis. The largest value throughout the volume in each permuted group level T-map is then taken to form the null distribution of t-values. A total of 1,000 permutations were analysed to generate the full null distribution. Give the alpha (=0.05 two tailed), if the t-value of the real data exceeds the 97.5 percentile or falls below the 2.5 percentile, it is considered significant.

### Stage Two: Analysis of Evoked Time Series at Selected Virtual Electrode Locations

#### Overview

Based on the results of the beamformer metric, the time series of local maxima voxel that demonstrated significant power differences between conditions was extracted for each condition. This was achieved by passing sensory data for each trial in each condition back through the beamformer weights at the specified brain location, in order to estimate a 3D time series referred to as a virtual electrode (VE). Based on the orientation that maximises the difference between conditions at this location, each trial time series could be expressed as a one-dimensional vector. This time series could then be dealt with in a similar way to normally epoched data. For each participant, epochs of the time series of every trial in each condition were extract from the identified VEs, these could then be used in statistical comparisons. First comparisons are performed for window selection, to identify a time window in which there were significant differences between conditions. These comparisons were performed only between the condition associated with the beamformer metric localisation. Then, in the average amplitude section of the VE analysis a summary statistic, across the selected window, was extracted from the timeseries of each condition. VEs from the Identity localisation could then be tested for a double dissociation against Orientation conditions within this time window, and similar VE from the Orientation localisation could be tested for double dissociation against Identity conditions.

#### Window Selection

To define temporal windows of interest for further analysis, an initial analysis of time series data for a subset VE identified in the beamformer metric analysis used timepoint by timepoint t-tests (0-500ms). This investigated difference between conditions, separately for the Identity VEs in the Identity comparison (CCIPO; CCIVO < VIPO; VIVO), and the Orientation VEs in the Orientation comparisons (CCIPO; VIPO; PIPO < CCIVO; VIVO; PIVO). Subset selection was based on proximity to other VEs and the intensity of beamformer metric t-values (i.e. for VEs less than 20mm^3^ from any other VE maxima, the VE with the highest beamformer metric t-intensity was selected). To correct for multiple comparisons in these t-test, the analysis used a temporal clustering algorithm. An initial common height threshold was selected (one-tailed *p* < .05) for t-values (*t* = 1.721) based on the sample size (*df* =21). This criterion was then used to form temporal clusters of t-values that reached this threshold for each VE. Each t-value in the temporal cluster had the t-value threshold subtracted from it. The remaining above threshold t-values were then summed, to give a value for the entire cluster.

To determine whether a given cluster was significant it was then compared to a null distribution of *t*-values. The null distribution was computed using a sign flip permutation of difference waves for the trials of specific comparisons. This procedure produces a much more realistic null than comparing t-values to zero. The sign (+/−) was randomly flipped (10,000 permutation) for participants difference waveforms, so that any condition effect should be destroyed. For each permutation, the temporal clustering analysis applied to the real data was repeated using the same minimum height threshold (*t* = 1.721) on time point by time point t-tests to form temporal clusters. The largest cluster value in each permutation then contributed to the null distribution. Each considered all VEs in the comparison together so as to control for comparisons across multiple VEs. The *t*-value for significance (one-tailed) in the real t-values was determined by the value in the 95^th^ percentile of this null distribution. Within each VE Clusters that met this threshold criterion were considered significant and used in further analysis.

#### Average Amplitude Analysis

Based on VEs and their respective time windows identified by the temporal clustering permutation test for each comparison (Identity or Orientation). The average amplitude within each of the four condition (Violated Orientation, Non-violated Orientation, Violated Identity and Non-violated Identity), across the specified time window was extracted within the respective VE. These values could then be analysed using an ANOVA in each VE with the factors Type of Expectation (Orientation vs. Identity) and Predictability (Violated vs. Non-Violated). Crucially, a significant interaction of these factors in any time window identifies a dissociation between Orientation conditions and Identity conditions.

## Results

### Stage One: Localising Evoked Responses in Violated Identities and Violated Orientations using the Evoked Power Variance Metric in Spatial Beamformer Analysis

The initial analysis considered the whole brain using the beamformer metric analysis described above. After an initial mask was applied to the data so that white matter was excluded from the analysis, this approach enabled the localisation of brain regions activated in the identity localisations and the orientation localisations separately. To provide sufficient power to localisations, the analysis of each stimulus attribute compared any Non-violation and Violation conditions that were entirely matched transitions across this comparison. As such, the orientation localisations contained all six conditions (CCIPO; PIPO; VIPO vs. CCIVO; PIVO; VIVO, whereas the Identity localisations only considered four conditions (CCIPO; CCIVO vs. VIPO; VIVO), because PI conditions had no identical match in the VI condition. These localisations had notably different voxels of significance, following null label permutation significance thresholding, for the Orientation localisations and the Identity localisations. The Orientation localisation demonstrates significant differences in activity between PO and VO conditions in areas known to be involved in perception of motion such as the left hMT/V5 and left angular gyrus (Figure 2). By comparison, in the Identity localisation, differences between CCI and VI conditions were significant at the right temporal occipital fusiform, known to respond to changes in face identity (Figure 3).

**Figure 2.**
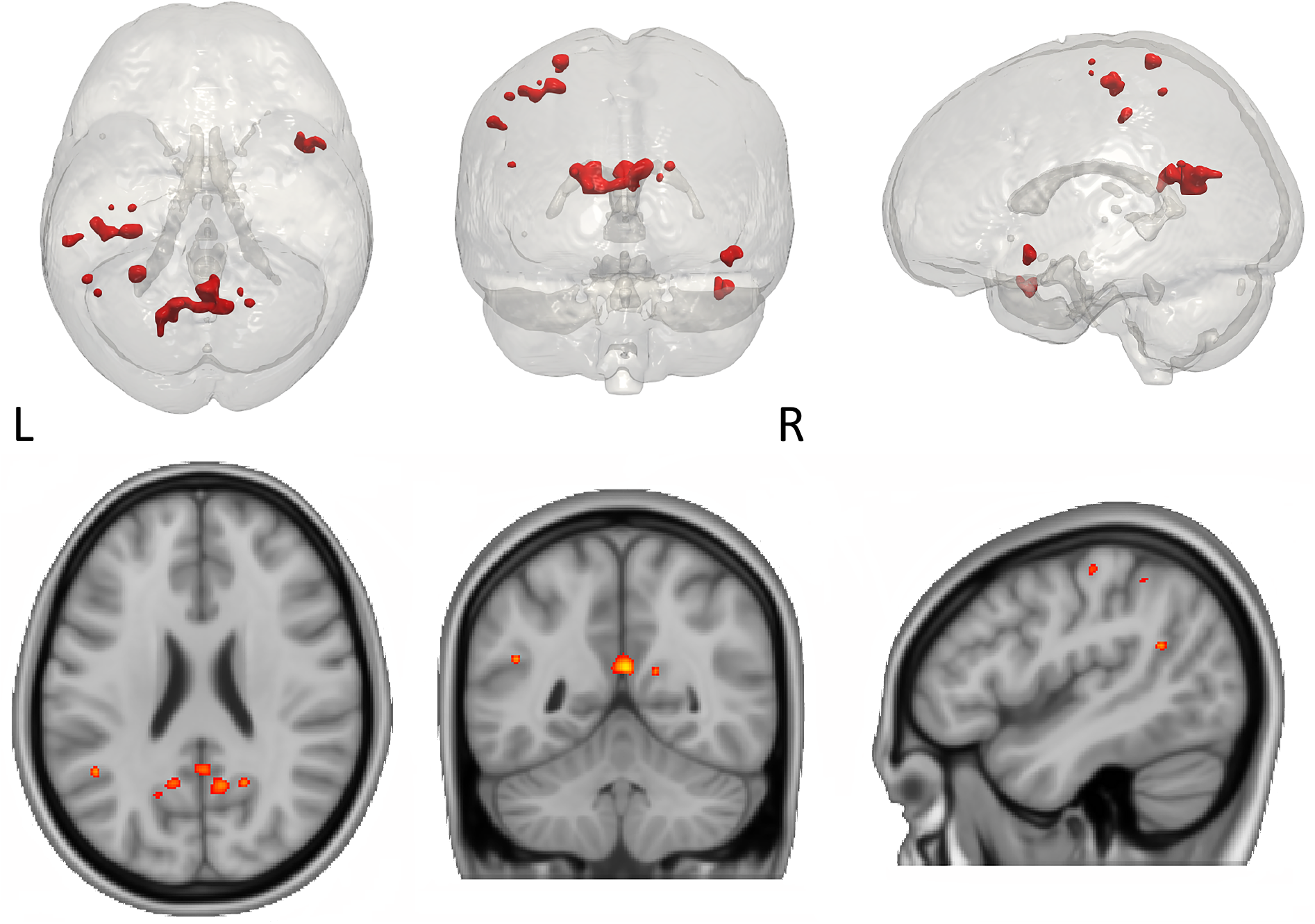
The bottom panel contains key selected slices of a t-values heat map from the Orientation localisation (CCIPO-PIPO-VIPO vs. CCIVO-PIVO-VIVO) displayed on the MNI 152 standard brain. The top panel contains a 3-D rendering of this same data on a transparant MNI brain with significant voxels displayed in red. 3-D meshes were constructed using ITK-SNAP (v.3.6.0) (Yushkevich et al., 2006)Thresholded to *p* <.05 (t = −6.451) based on a non-parametric null distribution created by permuting the conditions labels to produce a threshold for each localisation.

**Figure 3.**
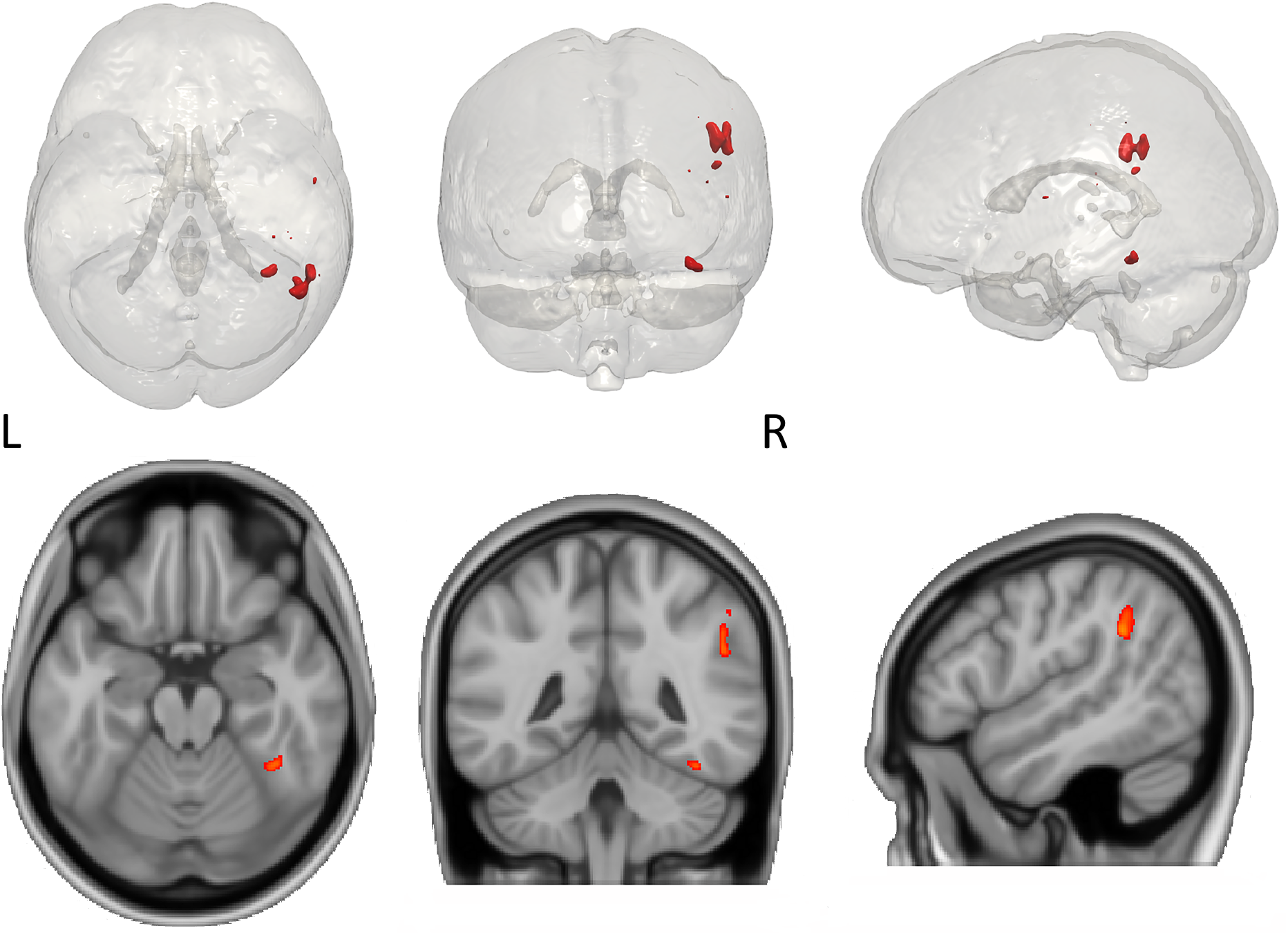
The bottom panel contains key selected slices of a t-values heatmap from the Identity localisation (CCIPO-CCIVO vs. VIPO-VIVO) displayed on the MNI 152 standard brain. The top panel contains a 3-D rendering of this same data on a transparant MNI brain with significant voxels displayed in red. 3-D meshes were constructed using ITK-SNAP (v.3.6.0) (Yushkevich et al., 2006). Thresholded to *p* <.05 (t = −6.648) based on a non-parametric null distribution created by permuting the conditions labels to produce a threshold for each localisation.

The beamformer metric *t*-values of localisations for Identity and Orientation conditions identified a number of peak local maxima that met the threshold for significance (see Table 1). It is important to note that the intensity values in the *t*-test localisations represented interpolated values as part of the transformation from the 5mm × 5mm to the MNI 152 standardised 1mm × 1mm voxel T1 brain image. Each significant local maxima was considered for subsequent VE analysis, where a comparison of the time series was performed for both the Identity and Orientation comparisons separately.

**Table 1.** Significant local maxima identified by the beamformer metric t-test for each experimental localisation.

| Beamformer Metric Localisation | Regions | Intensity (t) | x | y | z |
| --- | --- | --- | --- | --- | --- |
| Orientation | Posterior cingulate cortex | -9.969 | 3 | -51 | 19 |
|  | Left Precuneus | -9.514 | -18 | -62 | 19 |
|  | Left Postcentral gyrus | -9.212 | -26 | -41 | 67 |
|  | Right Precuneus | -8.555 | 9 | -58 | 22 |
|  | Left Angular gyrus | -8.555 | -45 | -52 | 23 |
|  | Right Precuneus | -8.303 | 19 | -57 | 22 |
|  | Right Temporal pole | -7.950 | 49 | 12 | -31 |
|  | Right Temporal pole | -7.899 | 47 | 13 | -17 |
|  | Left Supramarginal gyrus | -7.899 | -56 | -30 | 43 |
|  | Left Postcentral gyrus | -7.849 | -43 | -22 | 56 |
|  | Right Temporal pole | -7.395 | 53 | 13 | -15 |
|  | Left Supramarginal gyrus | -7.142 | -64 | -28 | 31 |
|  | Right Supramarginal gyrus | -7.142 | 57 | -31 | 38 |
|  | Left hMT/V5 | -6.739 | -43 | -68 | -2 |
|  | Right Superior Frontal gyrus | -6.739 | 21 | 15 | 49 |
| Identity | Right Angular gyrus | -9.464 | 46 | -52 | 33 |
|  | Right Angular gyrus | -8.257 | 51 | -43 | 30 |
|  | Right Temporal Occipital Fusiform gyrus | -7.871 | 36 | -47 | -19 |
|  | Right Supramarginal gyrus | -6.953 | 41 | -27 | 44 |
|  | Right Central Opercular cortex | -6.857 | 55 | -6 | 10 |
|  | Right Parietal operculum | -6.857 | 37 | -30 | 20 |
|  | Right Planum temporal | -6.760 | 45 | -30 | 15 |

Note. Voxels are considered significant based on a non-parametric null distribution created by permuting the conditions labels to produce a threshold for each comparison (alpha = .05). Intensity values are based on *t*-tests of leave-one-out distributions for each condition. Locations are based on MNI 152 (x,y,z) coordinates.

### Stage Two: Analysis of Evoked Time Series at Selected Virtual Electrode Locations

#### Window Selection

At this stage, a subset of the local maxima (Table 1) were selected for further analysis according to the procedure for proximal VE defined in Stage Two of the Statistical Analysis section. The VE analysis extracted the estimated time course for each trial at each of the identified voxel locations (Table 2) for each of the four compound conditions: 1. CCIPO, PIPO, VIPO; 2. CCIVO, PIVO, VIVO; 3. CCIPO, CCIVO; 4. VIPO, VIVO. An ERF wave form could then be computed for trial time series generated at each VE, which could then be averaged to demonstrate the estimated response across this condition in each participant. Of the 15 VEs identified in the Orientation localisation analysis (see Figure 4-1 & Figure 4-2), 10 were selected for further analysis by extracting VEs. From the seven brain locations identified in the Identity localisation (see Figure 5-1 & Figure 5-2), five were included in the subsequent VEs analysis.

**Figure 4.**
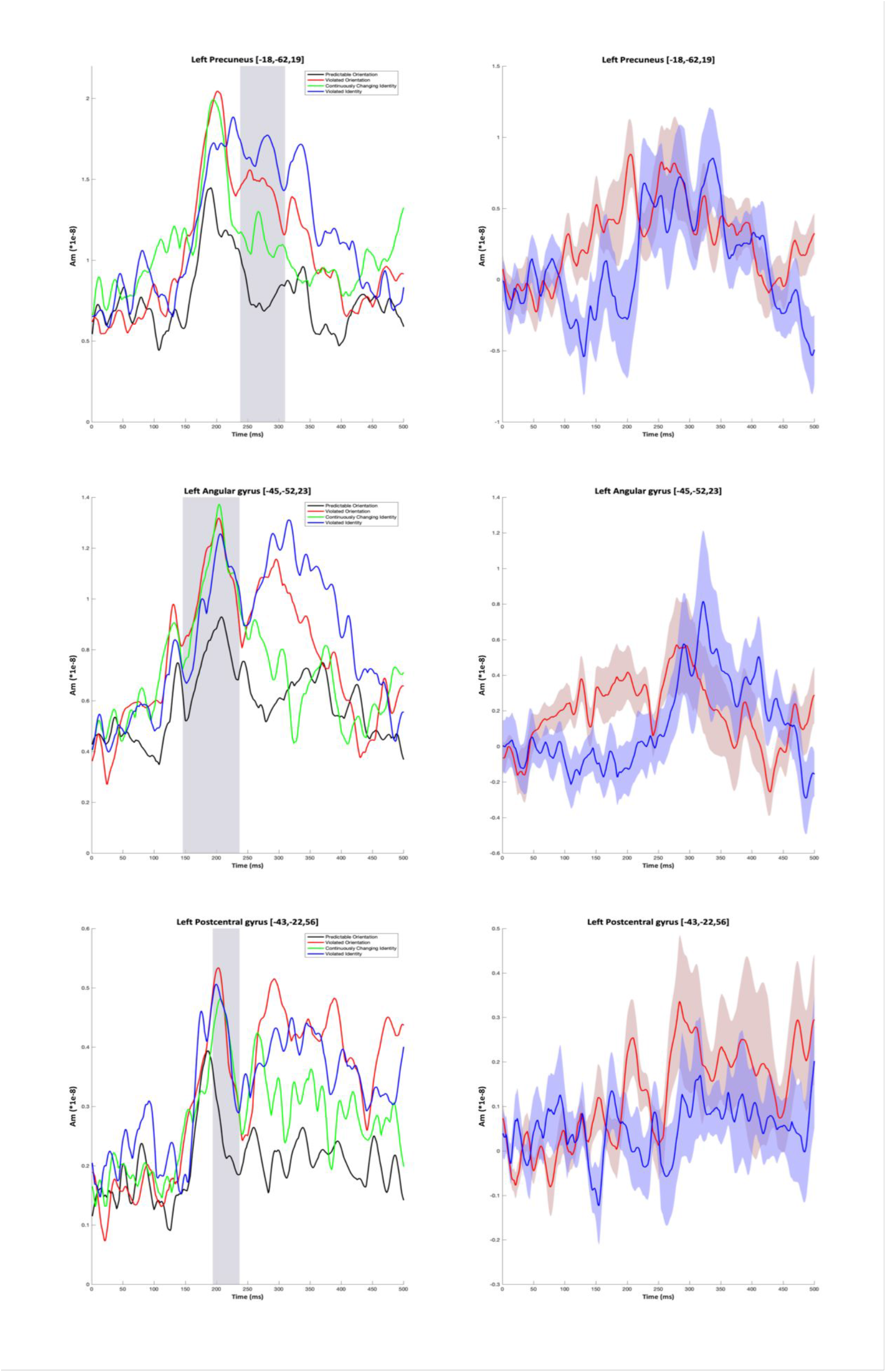
Selected VE time series from the Orientation window selection. On the left timeseries are presented for Non-violated Orientation (in black), Violated Orientation (in red), Non-violated Identity (in green), and Violated Identity conditions (in blue). On the right timeseries of the difference waveforms are presented for Orientation (in red) and Identity (in blue). Time series are baselined to the first 50ms after stimulus onset. Shaded grey sections on the left represent significant time windows for VE. Shading around timeseries on the right represent +/− 1 standard error for the respective time series.

**Figure 4-1.**
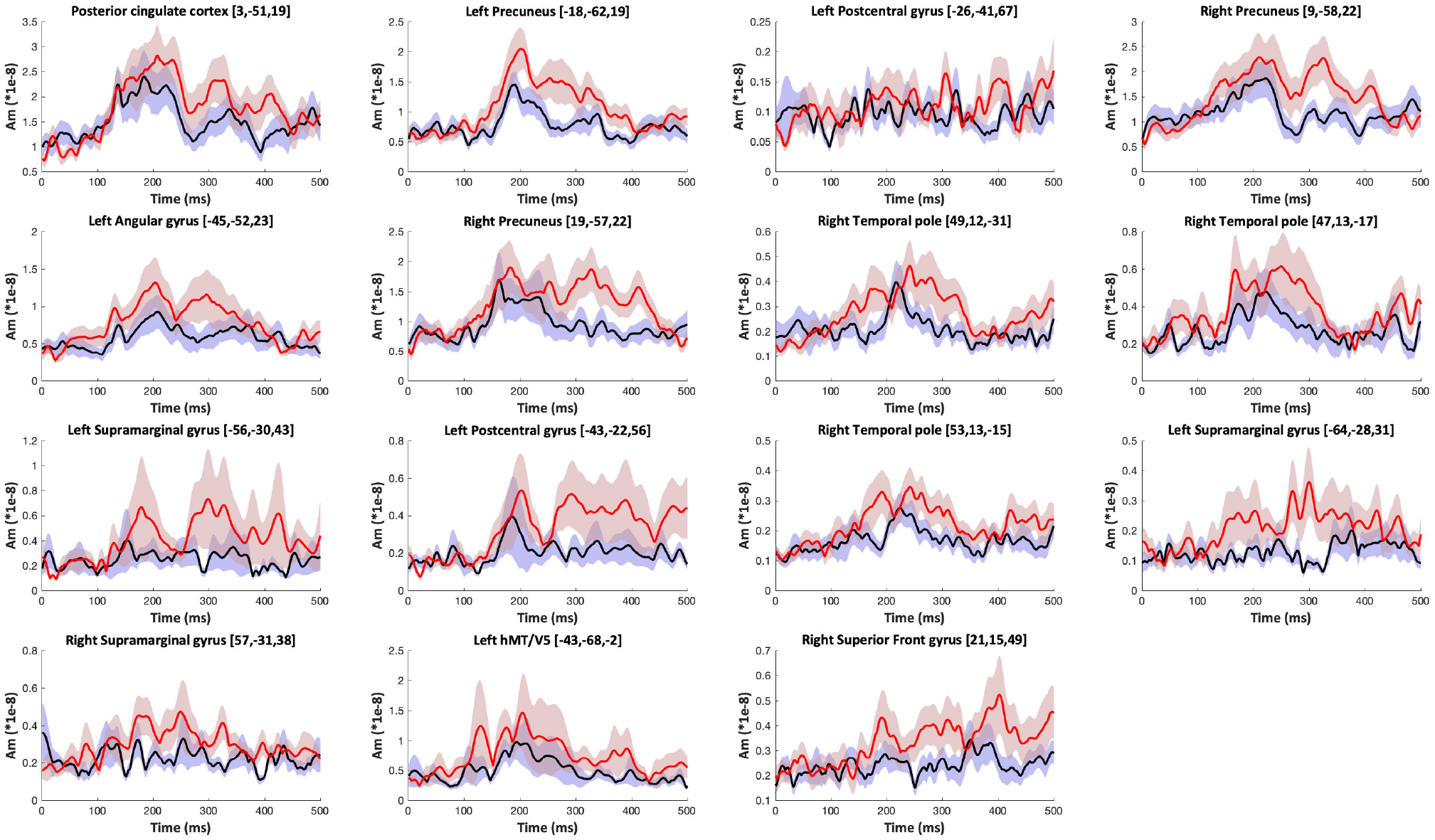
ERF time series for each of the voxels defined by the beamformer metric localisations of the Orientation comparison, with time series for the Orientation conditions in each VE plotted separately Time series run from the onset of the final stimulus to 500ms post stimulus and are displayed in absolute values. Time series are baselined to the first 50ms after stimulus onset. Black time series lines denote Non-violation conditions, whilst red lines denote Violation conditions. Shading around each time series denotes +/− 1 standard error.

**Figure 4-2.**
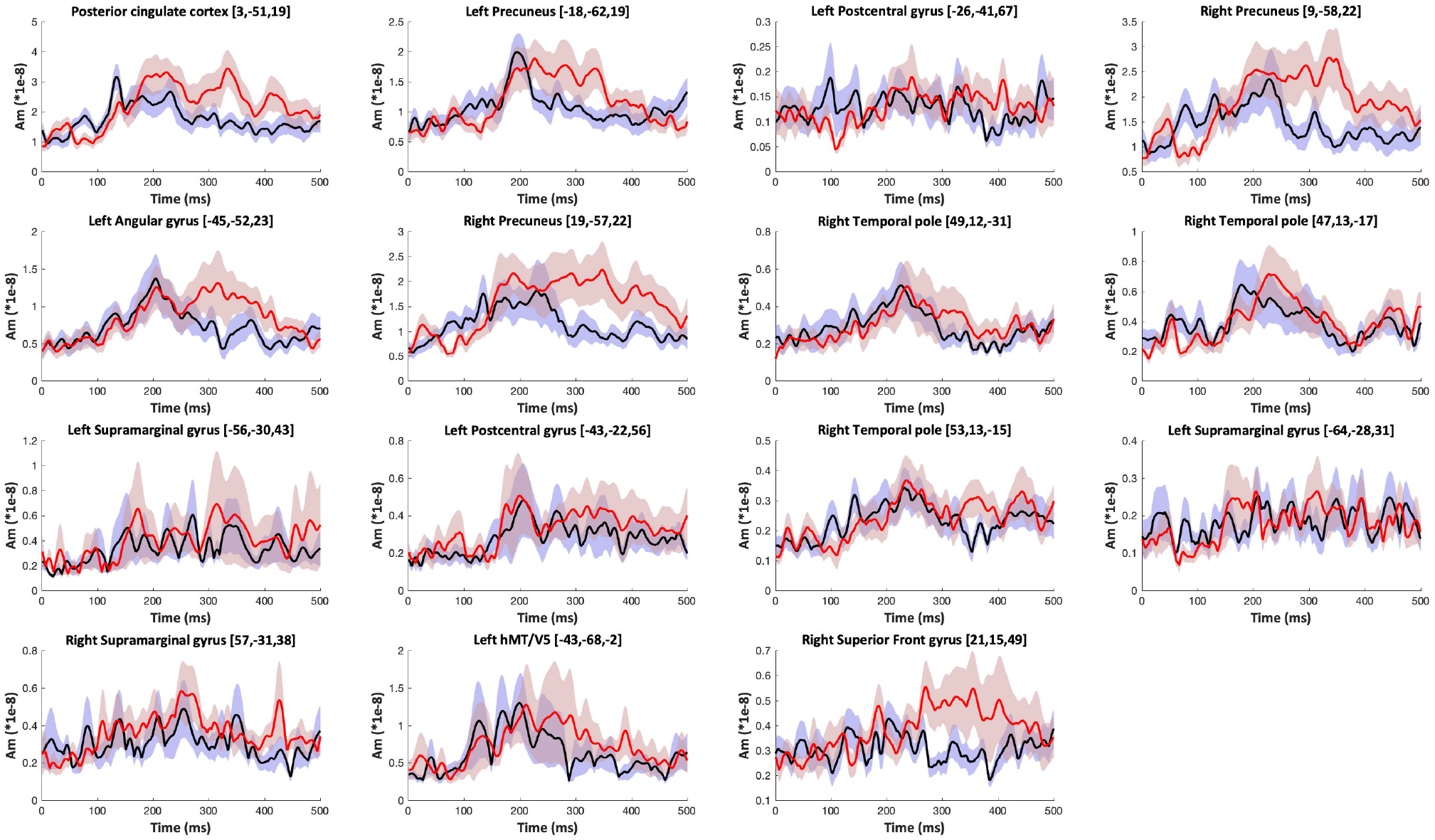
ERF time series for each of the voxels defined by the beamformer metric localisations of the Orientation comparison, with time series for the Identity conditions in each VE plotted separately. Time series run from the onset of the final stimulus to 500ms post stimulus and are displayed in absolute values. Time series are baselined to the first 50ms after stimulus onset. Black time series lines denote Non-violation conditions, whilst red lines denote Violation conditions. Shading around each time series denotes +/− 1 standard error.

**Figure 5.**
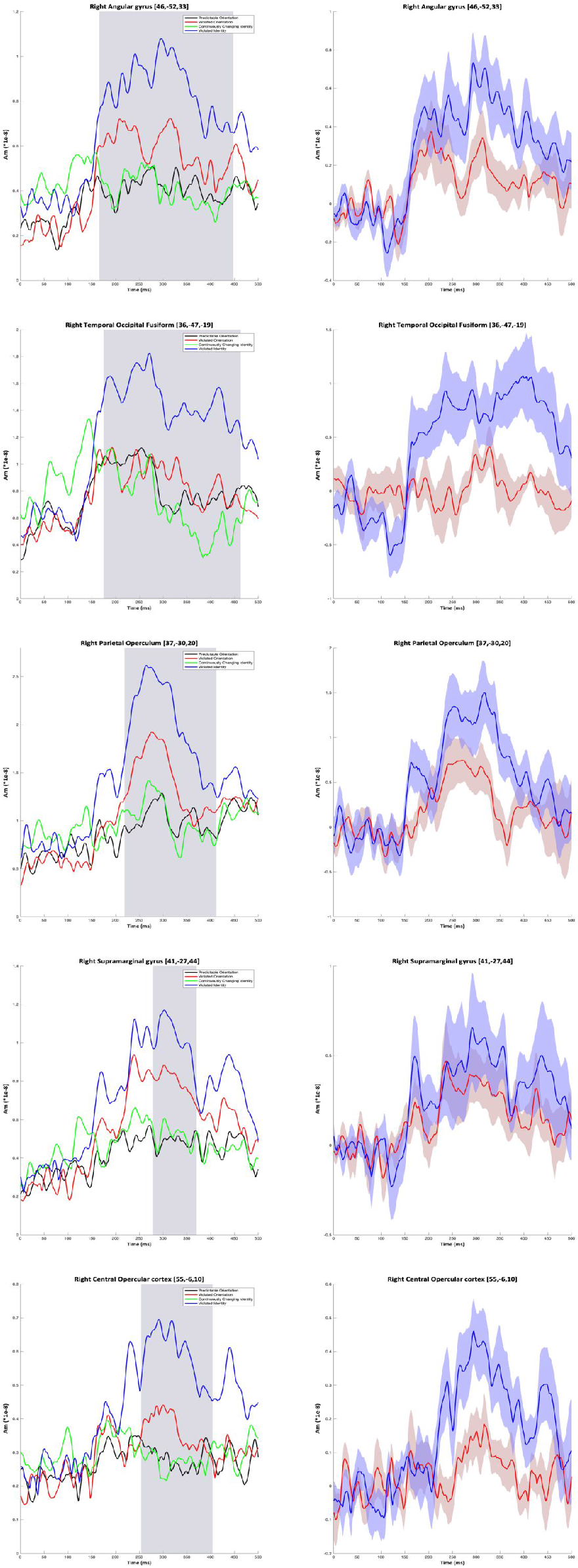
Selected VE time series from the Identity window selection. On the left timeseries are presented for Non-violated Orientation (in black), Violated Orientation (in red), Non-violated Identity (in green), and Violated Identity conditions (in blue). On the right timeseries of the difference waveforms are presented for Orientation (in red) and Identity (in blue). Time series are baselined to the first 50ms after stimulus onset. Shaded grey sections on the left represent significant time windows for VE. Shading around timeseries on the right represent +/− 1 standard error for the respective time series.

**Figure 5-1.**
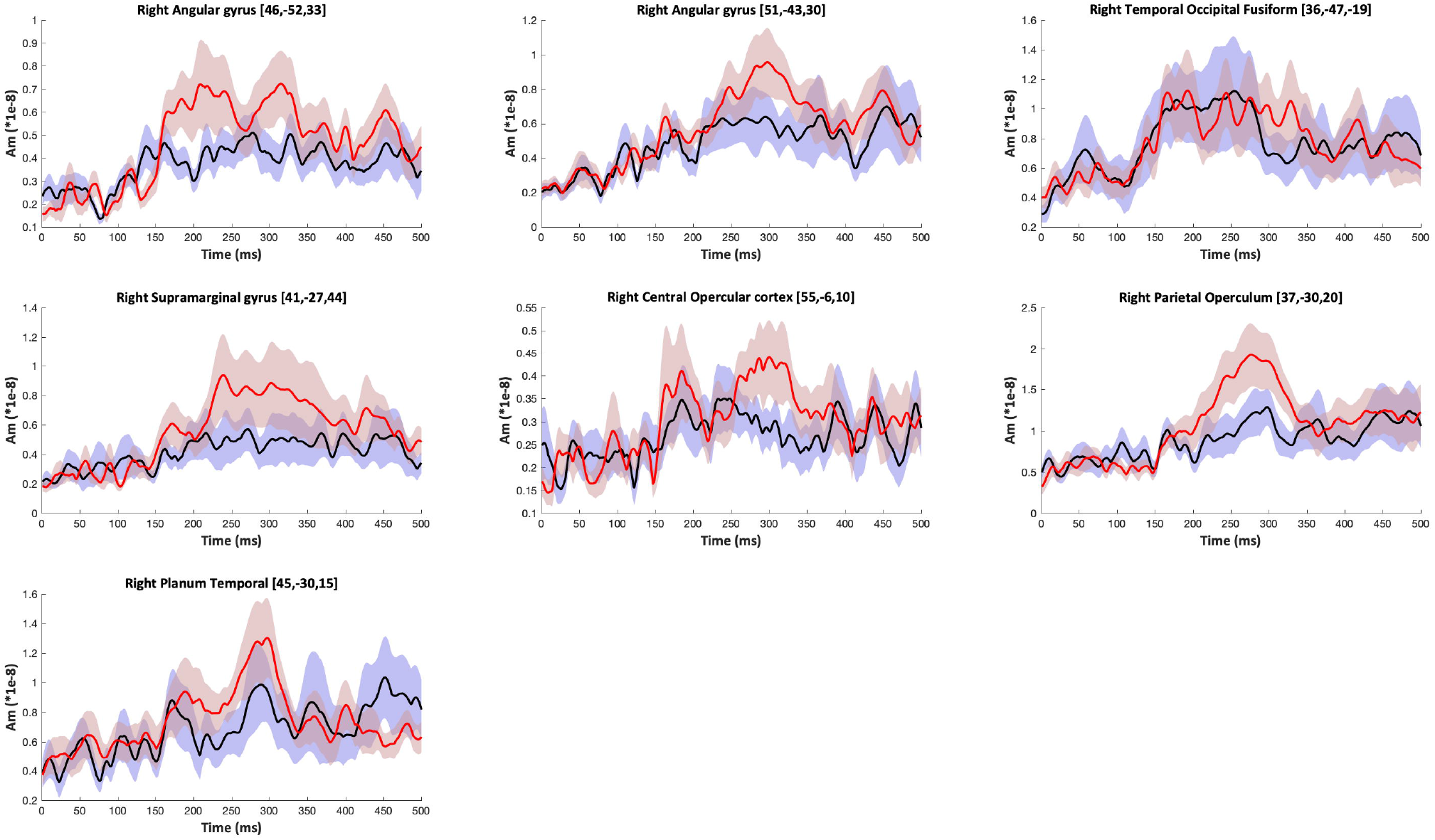
ERF time series for each of the voxels defined by the beamformer metric localisations of the Identity comparison, with time series for the Orientation conditions in each VE plotted separately Time series run from the onset of the final stimulus to 500ms post stimulus and are displayed in absolute values. Time series are baselined to the first 50ms after stimulus onset. Black time series lines denote Non-violation conditions, whilst red lines denote Violation conditions. Shading around each time series denotes +/− 1 standard error.

**Figure 5-2.**
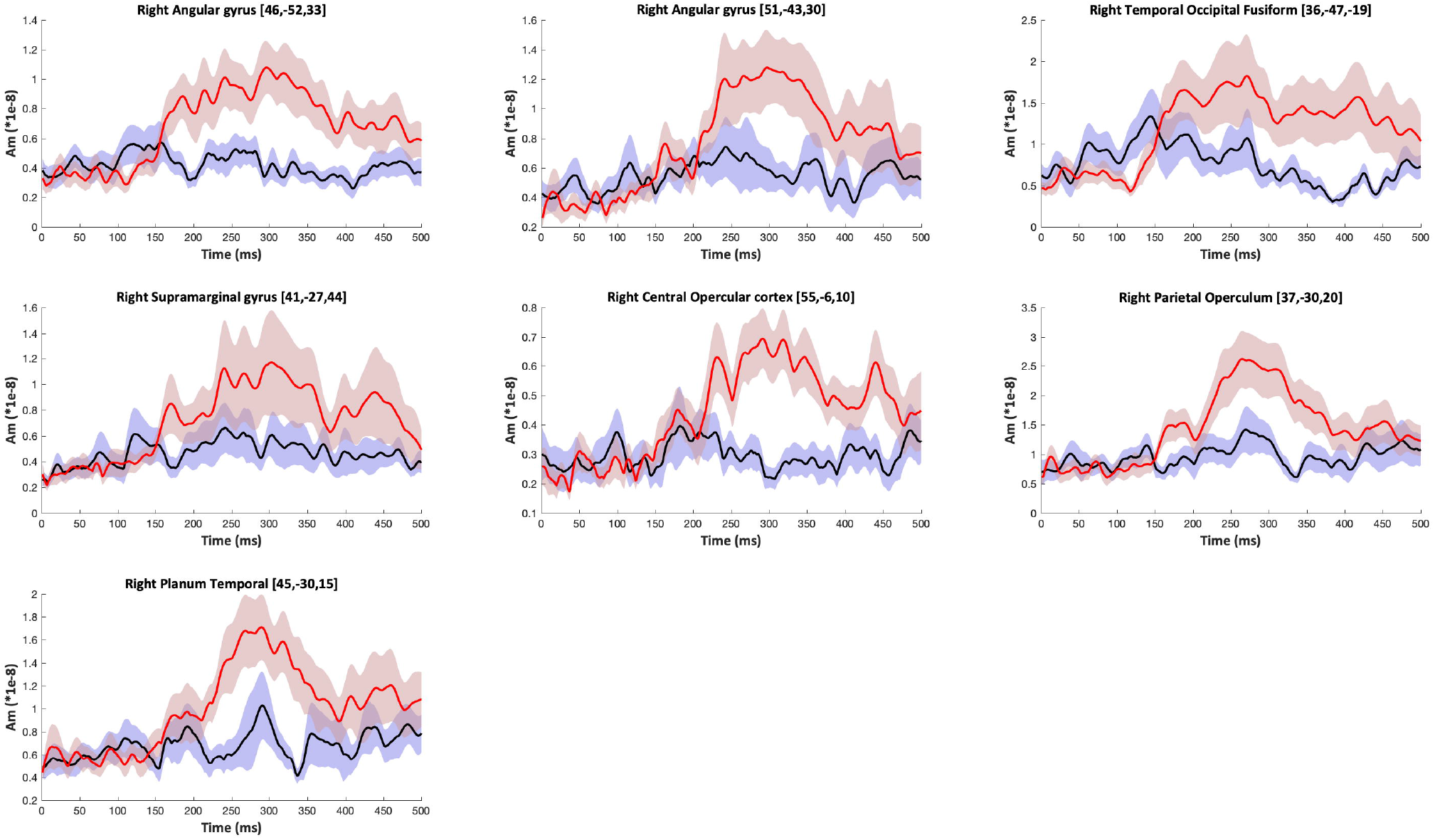
ERF time series for each of the voxels defined by the beamformer metric localisations of the Identity comparison, with time series for the Identity conditions in each VE plotted separately. Time series run from the onset of the final stimulus to 500ms post stimulus and are displayed in absolute values. Time series are baselined to the first 50ms after stimulus onset. Black time series lines denote Non-violation conditions, whilst red lines denote Violation conditions. Shading around each time series denotes +/− 1 standard error.

**Table 2.** VE selection for timepoint by timepoint t-test analysis

| Beamformer Metric Localisation | Index | Locations | x | y | z | Intensity (t) |
| --- | --- | --- | --- | --- | --- | --- |
| Orientation | 1 | Left hMT/V5 | -43 | -68 | -2 | -6.739 |
|  | 2 | Left Precuneus | -18 | -62 | 19 | -9.514 |
|  | 3 | Left Angular Gyrus | -45 | -52 | 23 | -8.555 |
|  | 4 | Posterior Cingulate | 3 | -51 | 19 | -9.969 |
|  | 5 | Left Postcentral Gyrus | -26 | -41 | 67 | -9.212 |
|  | 6 | Right Supramarginal Gyrus | 57 | -31 | 38 | -7.142 |
|  | 7 | Left Supramarginal Gyrus | -56 | -30 | 43 | -7.899 |
|  | 8 | Left Postcentral Gyrus | -43 | -22 | 56 | -7.849 |
|  | 9 | Right Temporal Pole | 49 | 12 | -31 | -7.950 |
|  | 10 | Right Superior Frontal Gyrus | 21 | 15 | 49 | -6.739 |
| Identity | 1 | Right Angular Gyrus | 46 | -52 | 33 | -9.464 |
|  | 2 | Right Temporal Occipital Fusiform | 36 | -47 | -19 | -7.871 |
|  | 3 | Right Parietal Operculum | 37 | -30 | 20 | -6.857 |
|  | 4 | Right Supramarginal Gyrus | 41 | -27 | 44 | -6.953 |
|  | 5 | Right Central Opercular Cortex | 55 | -6 | 10 | -6.857 |

Note. This VE subset was reordered according to their y coordinates in MNI space. Indices are used in subsequent t-test to denote VEs.

To define the select the appropriate time windows and VEs for the average amplitude analysis. The comparisons used in the beamformer metric were repeated in the VE analysis, however, instead of collapsing across a time window to analyse total power, the VE analysis was performed at each time point of the time series. Accordingly, two sets of cluster permutation t-tests were performed (see Figure 6-1). One set of tests was performed at the associated VEs locations comparing Orientation conditions (CCIPO, PIPO, VIPO < CCIVO, PIVO, VIVO). A second set of tests was performed on Identity VEs comparing Identity conditions (CCIPO, CCIVO < VIPO, VIVO). Only VEs that contained significant clusters were included in the final level of the analysis. Two null distributions were created for the analysis, one for comparisons over Orientation VEs and another for comparisons in identity VEs. This revealed a number of clusters that surpassed the significance threshold value in the given comparison (see Table 3). The orientation comparisons contributed three VEs (Figure 4) that surpassed a threshold value of 45.14 (*p* < .05) for final analysis, whilst five VEs from the Identity comparison (Figure 5) exceeded the permuted threshold value of 62.48 (*p* < .05). Significant Temporal cluster were identified for further analysis in each comparison within both early and midlatency time-windows.

**Figure 6.**
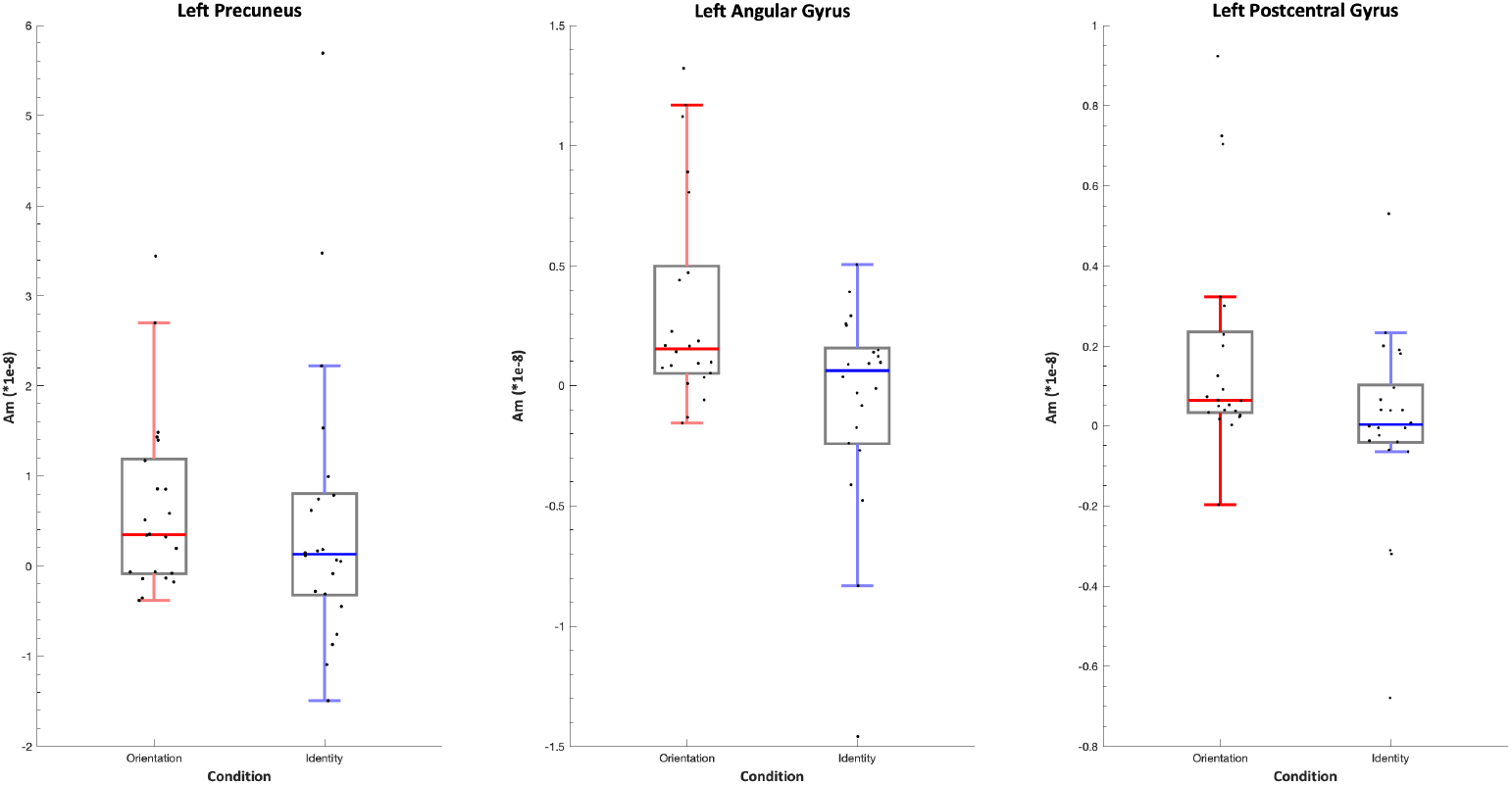
Difference box plot of Identity and Orientation condition (Violated – Non-violated) for the average amplitude across selected windows from the Orientation comparison in specified VE time series (see Table 3). Mid line represents data median and the box represent the upper and lower quartiles of the data. Whiskers represent data extremes and data points are plotted in black.

**Figure 6-1.**
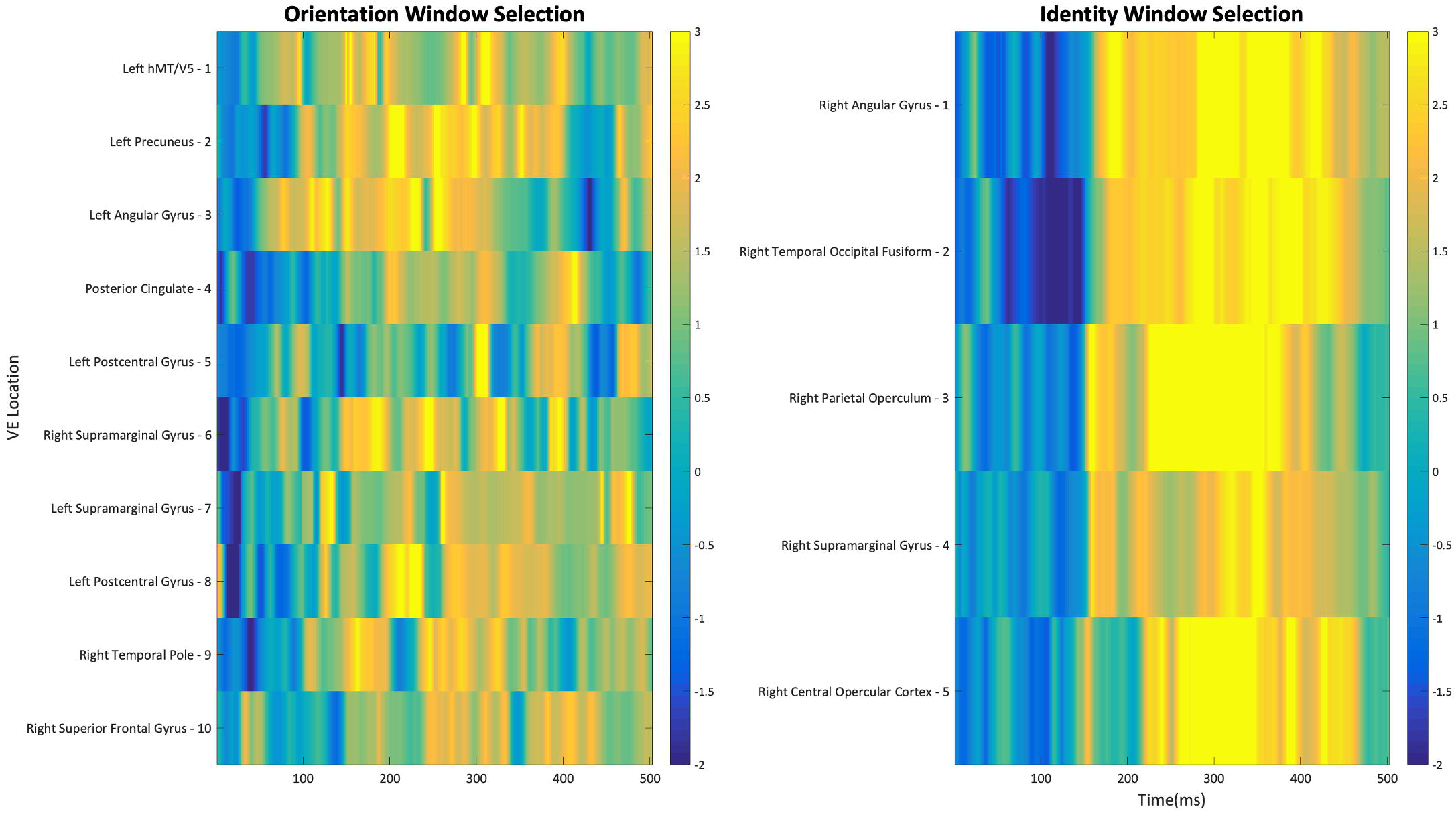
A *t*-test matrix for the VEs identified in each beamformer metric localisation on the y-axis (as listed in Table 2), with timepoints (0-500ms) on the x-axis. Orientation comparisons refer to *t*-tests between Orientation conditions (as in Table 2), whilst Identity comparisons refer to *t*-tests between Identity conditions (as Table 2). Each matrix uses a standardised *t*-threshold minimum (−2) and maximum (3).

**Table 3.**
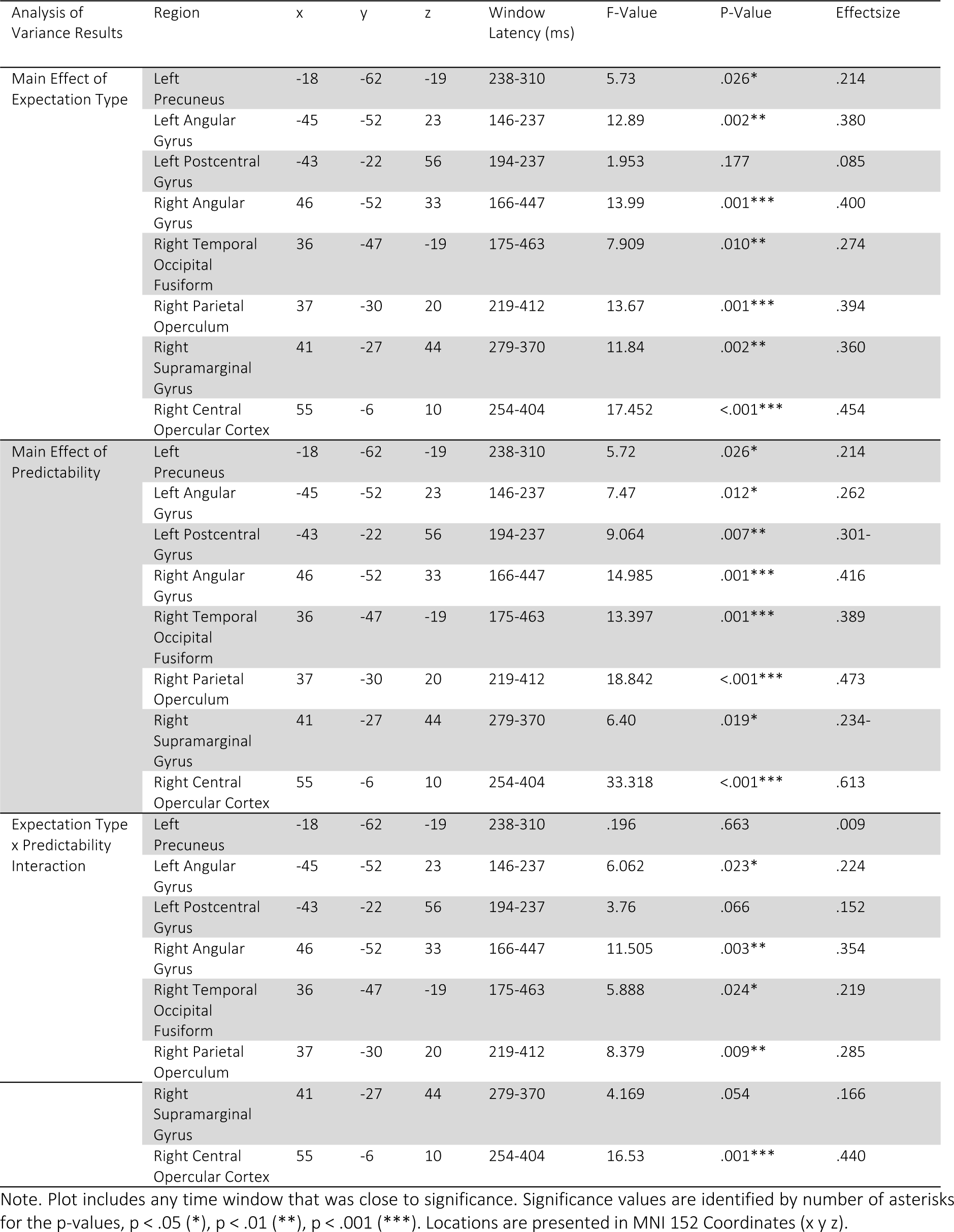
List ANOVA comparisons for VE comparison based on the averaged amplitude of significant temporal clusters.

#### Average Amplitude Analysis

To quantify these differences an ANOVA was performed between types of expectation (Orientation vs. Identity) and predictability (Violated vs. Non-violated). The results (table 5) highlight some regions that demonstrate greater sensitivity to some Orientation conditions than Identity and vice versa for others. In the early-latency window selected (146-237ms) for the Left Angular Gyrus VE (see Figure 4) consistent higher response activity can be seen to Violated Orientation conditions as compared to Non-violated Orientation, whilst there a less consistent difference in these VEs between identity conditions.

By comparison of the selected time window (175-463ms) in the VE located in the FG (Figure 5) Violated Identity conditions demonstrate consistently larger responses as compared to Non-violated Identity Conditions, not present to orientation conditions. The interaction in this comparison was fundamental to identifying any double dissociations in the present study. Importantly, as suggested by the timeseries data the left angular gyrus and fusiform gyrus VEs demonstrated a clear interaction. Differences boxplots between orientation and identity conditions in Orientation VEs (Figure 6) and Identity VEs (Figure 7) highlight the presence of a double dissociation for each of these sites.

**Figure 7.**
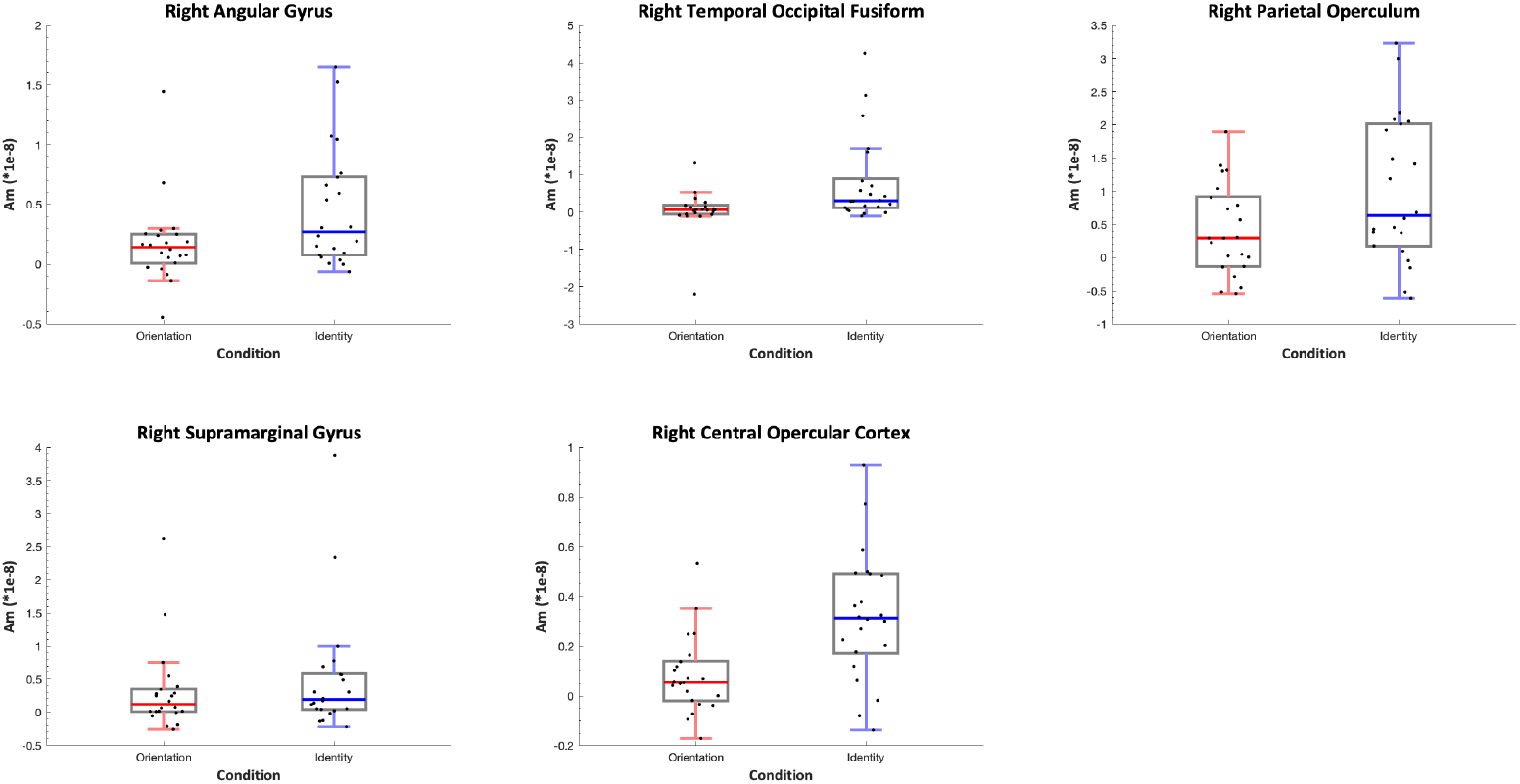
Difference box plot of Identity and Orientation condition (Violated – Non-violated) for the average amplitude across selected windows from the Identity comparison in specified VE time series (see Table 3). Mid line represents data median and the box represent the upper and lower quartiles of the data. Whiskers represent data extremes and data points are plotted in black.

## Discussion

Our study explored how dorsal and ventral visual stream areas are involved in processing early latency prediction-error detection. We designed our study to build upon the paradigm used by Johnston et al. (2017) which identified early-mid latency EEG/MEG prediction-error signals in the dorsal visual stream to violated expectations about stimulus orientation/trajectory, with localisations consistent with cortical areas processing motion and orientation. Here, by comparing activity related to violations of orientation with activity related to identity, we extend our previous findings by identifying and dissociating predictive processes in both dorsal and ventral visual streams.

MEG Beamformer contrasts revealed a set of cortical regions that respectively showed evidence of differences in activation to violations of expectations about head orientation (including regions of the dorsal and medial parieto-temporal cortices, and the left temporal pole) and to violations of expectation with respect to person identity (including the fusiform gyrus, and regions of right dorsal parieto-temporal and opercular cortices). The novel beamformer metric used here for whole brain analysis necessitates a reasonably long window in order to establish stable measures of covariance (in this case we used a 60-500ms post stimulus onset since this period encompassed the majority of the evoked power in the signal). This metric is sensitive to differences in evoked power between pairs of conditions across that time period. We estimated Virtual Electrode timeseries based on the ROIs identified by the beamformer metric, and a used temporal clustering permutation method to establish the time periods following stimulus onset during which the timeseries to Non-violated and Violated expectation trials showed significantly different amplitudes. This revealed a subset of ROIS that showed temporally constrained differences in signal amplitude across conditions: the left precuneus; left angular gyrus; left postcentral gyrus; right angular gyrus; right temporal occipital fusiform; right parietal operculum; right supramarginal gyrus and right central opercular cortex.

That the other ROIs identified by the beamformer analysis did not show significant differences between Non-violated and Violated Orientations or Identities *across specific time-windows* may reflect either insufficient power to detect such differences, or, that the consistent differences in evoked power between the conditions was not time-locked to a specific latency window. For VEs where there was evidence of temporally constrained expectation violation signals, we extracted averaged signal amplitudes for those time windows for all 4 types of trial (Predictable Orientation, Unpredictable Orientation, Predictable Identity, Unpredictable Identity) and compared signal amplitudes using ANOVAs. This revealed a pattern of findings consistent with evidence of early-mid latency double-dissociations across a number of cortical regions. Specifically, both the left Angular Gyrus (146-237ms) and the left Postcentral Gyrus (194-237ms) showed evidence of greater activation to violated expectations relating to head orientation than to violated expectations about face identity. By contrast, the right Fusiform Gyrus (175-463ms), right Parietal (219-412) and Central Opercular (254-404) cortices showed evidence of greater activation to violated expectations about face identity than to violated expectations about head orientation.

These findings are generally consistent with our predictions. Dorsal visual stream area, the left Angular gyrus has previously been implicated in visuo-spatial tasks, such as directionality discrimination (Ardila et al., 2000; Hirnstein et al., 2011) and number line representations (Cattaneo et al., 2009). In the current study, the angular gyrus showed evidence of generating expectation violation signals to unexpected head orientations but not to unexpected face identities. This is consistent with the core idea that prediction-errors about orientation are detected at the level of the processing hierarchy that resolves object orientation. This result is also consistent with our previous research localising prediction-error signals to face and body orientation (Johnston et al., 2017) since the (right) angular gyrus was one of the main ROIs reported in that study.

In the current study there were also some inconsistencies with our previous study, which identified largely right lateralised (temporo-parietal) sources, whereas for orientation violation signals in the current study sources were predominantly left lateralised. It is not clear why these inconsistencies were present, however, there were a number of important differences between the two experiments including. The previous study employed an MEG scanner using only magnetometers sensors, whilst in the current study the MEG system used a combination of both magnetometers sensors and gradiometer sensors. These different types of sensors have differential sensitivity with respect to detecting sources depending on the depth and orientation of the sources magnetic field. Furthermore, the previous study used different stimulus sequences (i.e. bodies as well as head), which may have played a role. Moreover, these same right-lateralised temporo-parietal sources do show evidence of activations to orientational violations in the current study, which do not meet the criteria for statistical significance. Significant responses to orientation in the left postcentral gyrus demonstrate localisations consistent with the head and face area of the somatosensory cortex. These findings could be reflective of error checking processes in models of head and face orientation.

A number of cortical areas showed evidence of greater early-mid latency violations to Identity violation than to violated Orientation. Our findings in the FG are consistent with fMRI adaptation data demonstrating a reduction in responses after repeated identities (Andrews and Ewbank, 2004), not present when time variant aspects of faces were changed. Our findings are also consistent with work by Simpson et al. (2015) showing adaptation effects to repeated identities in the FG despite not inducing strong identity expectation. The findings are also consistent our previous study (Johnston et al., 2016) showing greater activity at early latencies to rare identities than frequent identities in ambient image presentations. A right lateralisation of this response, as in the present study, is not uncommon in response to the presentation of face stimuli (Kanwisher et al., 1997; Morris et al., 2007).

The opercular cortex regions showed greater surprise to identity violation than to orientation. This is perhaps surprising as the opercular cortices are not within the ventral stream. Indeed, Johnston et al (2017) previously reported ROIs in right parietal opercular cortex and right central opercular cortex to violations of expectation to face and body orientation. In the present study we found a greater prediction-error signal to identity violations in these regions than to Orientation Violations. These results might initially seem at odds with our previous reported results, however, a careful examination of the data suggests otherwise (Figure 5-1). The current data suggest differences between Violated and Non-violated orientation are present but that the magnitude of violation is much bigger to identity.

The findings of Robinson et al. (2018) support this perspective as they demonstrate that the magnitude of prediction-errors is surprise dose dependent. Given that face stimuli are highly salient (Langton et al., 2008), strong models of identity will create more surprise when violated, as compared to a less salient stimulus feature such as head orientation, where changes in orientation are much more likely. The current study clearly demonstrates prediction-error signalling to violated expectations in both head orientation and person identity. These data cannot be attributed to low-level difference between stimuli, because the final sequence transition for trials in each expectation type (across which all comparisons were performed) was identically matched across Violated and Non-violated conditions. Therefore, any conditions differences after the onset of the final stimulus must be the result of prior expectation created across the preceding image sequence.

The study clearly identified a double dissociation of function between a ventral regions; the FG, which responds to violation in identity but not orientation and a dorsal region, and the left angular gyrus, which is sensitive to violation in orientation but not identity. The window identified for each of these VEs begun at similar early-latencies suggesting a common process for the perception of violations across different types of expectation within the perception of faces. Thus, we have shown for the first time, that prediction-error signals to specific stimulus attributes are localised to distinct cortical regions. These findings are consistent with the idea that the generation of prediction-error signals occurs at the hierarchical level where the visual feature is processed (Friston, 2005). In the present study we have demonstrated that prediction-error signal to different attributes of a well-studied type of stimulus category, the face, are generated in distinct cortical regions. We believe that this phenomenon is likely to generalise to different types of stimuli and stimulus attributes, and across different sensor and perceptual modalities.

## Acknowledgements

We acknowledge the Australian National Imaging Facility for the financial support of W.W. and the MEG system at Swinburne University of Technology. We gratefully acknowledge the gentle encouragement of the Oily Rag Foundation (EN10005). We’d also like to thank Jessica Guy, Deborah Loats and the rest of the SUT Babylab team for their involvement in data collection.

Author Contribution
J. E. R. and P. J. J. conceived the study, designed the study, and led writing of the manuscript. W. W. analysed the MEG data. S.L. and J. K. assisted in conception of the study and acquired the data. M. B. and A. W. Y. advised on the study design, data interpretation and writing up of the manuscript.

## Notes

**Conflict of Interest**: The authors declare no competing financial interests, in accordance with *JNeurosci’s* Policy on Conflict of Interest.

